# Approximated Gene Expression Trajectories (AGETs) for Gene Regulatory Network Inference on Cell Tracks

**DOI:** 10.1101/2022.01.12.476060

**Authors:** Kay Spiess, Shannon E. Taylor, Timothy Fulton, Kane Toh, Dillan Saunders, Seongwon Hwang, Yuxuan Wang, Brooks Paige, Benjamin Steventon, Berta Verd

## Abstract

The study of pattern formation has greatly benefited from our ability to reverse-engineer gene regulatory network (GRN) structure from spatio-temporal quantitative gene expression data. Traditional approaches omit tissue morphogenesis, and focus on systems where the timescales of pattern formation and morphogenesis can be separated. In such systems, pattern forms as an emergent property of the underlying GRN and mechanistic insight can be obtained from the GRNs alone. However, this is not the case in most animal patterning systems, where patterning and morphogenesis are co-occurring and tightly linked. To address the mechanisms driving pattern formation in such systems we need to adapt our GRN inference methodologies to explicitly accommodate cell movements and tissue shape changes. In this work we present a novel framework to reverse-engineer GRNs underlying pattern formation in tissues undergoing morphogenetic changes and cell rearrangements. By integrating quantitative data from live and fixed embryos, we approximate gene expression trajectories (AGETs) in single cells and use a subset to reverse-engineer candidate GRNs using a Markov Chain Monte Carlo approach. GRN fit is assessed by simulating on cell tracks (live-modelling) and comparing the output to quantitative data-sets. This framework generates candidate GRNs that recapitulate pattern formation at the level of the tissue and the single cell. To our knowledge, this inference methodology is the first to integrate cell movements and gene expression data, making it possible to reverse-engineer GRNs patterning tissues undergoing morphogenetic changes.

## Introduction

Embryonic pattern formation underlies much of the diversity of form observed in nature. As such, one of the main goals in developmental biology is to understand how spatio-temporal molecular patterns emerge in developing embryos, how they are maintained and how they can change over the course of evolution. Over the past three decades, the field has focused on the function and dynamics of the gene regulatory networks (GRNs) underlying these processes. GRNs can be formulated mathematically as non-linear systems of coupled differential equations whose parameters can be inferred from quantitative gene expression data: a methodology known as reverse-engineering (*1–8*). Reverse-engineering has been successfully applied to a myriad of systems, from the *Drosophila* blastoderm to the vertebrate neural tube (*9–12*), uncovering the mechanisms by which GRNs read out morphogen gradients (*12–17*), scale patterns (*18*), control the timing of differentiation (*19–21*), synchronise cellular fates (*22*) and evolve pattern formation (*23*).

In addition to the spatial patterning of gene expression within tissues, the developing embryo needs to deform and grow tissues to the correct shape and size during morphogenesis. Much of what we know about pattern formation has been learnt from reverse-engineering GRN structure in systems where the timescales of pattern formation and morphogenesis are different and can therefore be separated. In such systems, spatio-temporal gene expression profiles are typically obtained by measuring gene expression levels across the tissue of interest in fixed stained samples, and interpolating between measurements taken at different time points (*8*). The underlying and seldom stated assumption, is that the gene expression dynamics are much faster than the cell movements in the developing tissue, and that therefore cell movements can be ignored over the timescales at which the pattern forms. This is true in many systems and processes such as segmental patterning in early *Drosophila* embryogenesis. In systems where this is indeed the case, pattern formation can be considered an emergent property of GRN dynamics alone (*24*) and much insight can be drawn from analysing reverse-engineered GRNs (*10, 13*).

In systems where tissue patterning and tissue morphogenesis are coupled and occurring simultaneously, GRNs alone cannot account for the resulting patterns. This has been recently highlighted by work in organoids, where shape, size, and cell type distributions and proportions are difficult to control as a result of altered patterning due to abnormal morphogeneses in unconstrained tissue geometries (*25*). Therefore, in order to be able to understand developmental pattern formation in a broader range of systems, we have to address how morphogenesis and GRNs together control fate specification and embryonic organisation. Importantly, to be able to do this, we need novel reverse-engineering methodologies that will explicitly accommodate cell movements and tissue shape changes.

Here we present a methodology to reverse-engineer GRNs underlying pattern formation in tissues that are undergoing morphogenetic changes such as cell rearrangements. As a case study we focus on T-box gene patterning in the developing zebrafish presomitic mesoderm (PSM) (see schematic in Figure 1A and Supplementary Figure 1A). T-box genes coordinate fate specification along the PSM as cells move out of the tailbud and towards the somites (*26*). Cell movements in the PSM can be live-imaged and followed in 3D (*27*). By the time they reach a somite, cells in the PSM will have undergone a stereotypical progression of T-box gene expression: *tbxta* and *tbx16* in the tailbud, followed by *tbx16* in the posterior PSM and *tbx6* in the anterior PSM (Supplementary Figure 1A and D). The *tbx16*/*tbx6* boundary roughly marks the cells’ transition out of the tailbud and in zebrafish it is thought to correlate with marked changes in cell behaviours where extensive cell mixing in the tailbud gives way to reduced, almost nonexistent mixing and neighbourhood cohesion in the PSM (*28*). Therefore, while all cells will eventually have undergone the same gene expression progression, their expression dynamics will differ as cells spend variable amounts of time in the tailbud (*26*). Despite this, a tissue-level pattern forms which scales with PSM length over the course of posterior axial elongation and somitogenesis (*26*). T-box pattern formation in the developing zebrafish PSM is therefore a good example of a developmental process where the molecular pattern across the tissue is an emergent property of the GRN, the cell movements and tissue shape changes involved in the tissue’s morphogenesis.

**Figure 1:**
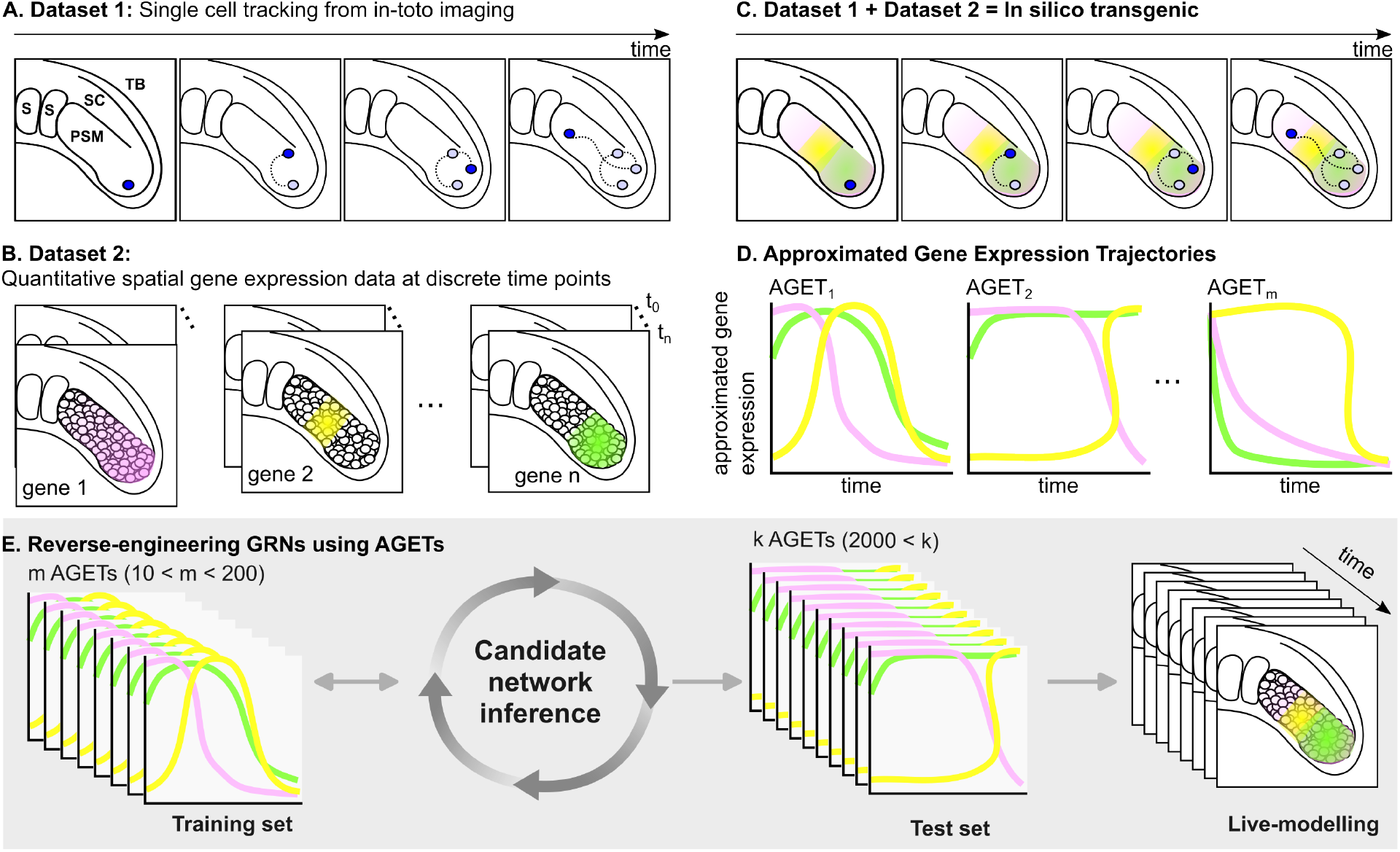
Approximated gene expression trajectories (AGETs) to reverse-engineer GRNs patterning tissues undergoing morphogenesis and cell rearrangements. The generation of AGETs requires the combination of two datasets: **A**. single cell tracking data and **B**. HCR gene expression data at the same stages as those covered by the tracking data. **C**. These two datasets are combined to generate and in silico transgenic reporter where every cell in the tracking data has been assigned a gene expression value for every time point using the HCR data. **E**. Subsets of AGETs can then be used to reverse-engineer GRNs, and candidate networks can be simulated using the initial and boundary conditions for all AGETs and represented back on the tracks to recapitulate pattern formation during tissue morphogenesis (live-modelling).

The reverse-engineering methodology presented in this paper accommodates cell movements and tissue shape changes, representing tissue morphogenesis explicitly when reverse-engineering GRNs. To do this, our methodology integrates two different kinds of quantitative data: cell tracking data obtained from live-imaging the developing tissue (Figure 1A) and three-dimensional quantitative gene expression of the genes and signalling pathways of interest over developmental time (Figure 1B). We project the 3D gene expression data onto the cell tracks to generate Approximated Gene Expression Trajectories (AGETs) in single cells (Figure 1C and D). A subset of AGETs is then used to reverse-engineer candidate GRNs applying a Markov Chain Monte Carlo (MCMC) approach (Figure 1E). The fit of the resulting candidate GRNs is assessed by simulating them in each cell in the tracks using initial and boundary conditions extracted directly from the gene expression data, a methodology that we refer to as “live-modelling”. The resulting well-fitting GRNs cluster, generating candidate GRNs that we further investigate and challenge using experimental work (*26*).

To our knowledge, this inference methodology is the first to integrate cell movements and gene expression data, making it possible to reverse-engineer the GRNs patterning tissues as they undergo morphogenesis. We hope that this toolbox will contribute to broaden the types of patterning systems that are studied quantitatively and mechanistically, increasing our understanding of pattern formation in development and evolution.

## Results and Discussion

### Approximating gene expression dynamics on single cell tracks: AGETs

The ideal data to reverse-engineer gene regulatory networks would be temporally accurate quantifications of gene expression dynamics at the single cell level as the tissue develops. Unfortunately, current state of the art in live gene expression reporter technology, while very advanced, cannot follow three genes and two signalling pathways simultaneously in space and time, while also ensuring that the dynamics of all reporters faithfully recapitulate the expression dynamics of the genes of interest. For this reason, it has been necessary to develop an alternative approach to effectively construct in-silico reporters based on approximating gene expression trajectories in the cells of the developing PSM, which we will from now on refer to as AGETs (Approximated Gene Expression Trajectories).

In brief, AGETs are obtained by projecting 3D spatial quantifications of gene expression in PSM cells obtained using HCRs and antibody stains, onto the cells present at each time frame of a time lapse movie of the developing PSM. The projected expression levels are used to assign gene and signalling expression levels in every cell in the time lapse. The result is an approximated gene expression trajectory for every cell in the time lapse, which can now be used to reverse-engineer gene regulatory networks which recapitulate T-box pattern formation on the developing PSM when simulated on the tracks.

### Data requirements and preparation

Two kinds of data are required to produce AGETs: cell tracks obtained from live-imaging the developing tissue of interest and quantitative spatial gene expression data at each developmental stage covered by the tracks.

In this case study, cell tracks were obtained by live-imaging a developing zebrafish tailbud with fluorescently labelled nuclei between the 22nd and 25th somite stages using a two-photon microscope (see (*27*) and Materials and Methods). Embryonic tailbuds were imaged for two hours at two-minute intervals, generating 61 consecutive frames. Each frame consists of a point cloud representing the position of single cells in 3D space. All the data were processed using a tracking algorithm in the image analysis software Imaris to obtain the position of single cells over time, and selected tracks were validated manually. The resulting data are a collection of cell tracks that describe how individual cells in the PSM move as the zebrafish tailbud develops.

A cell track provides spatial information over time but is devoid of any information regarding gene expression levels in each cell.

Gene expression levels were approximated from fixed tailbud samples stained for the T-box gene products using HCR (*29*) and antibody stains for the signals Wnt and FGF (see Materials and Methods). As the Tbox pattern scales with tissue growth over developmental time (*26*), embryos from different stages could be pooled together, however this method could also be adapted to patterning systems that do not scale with tissue growth. We stained for *tbxta, tbx16, tbx6*, and DAPI on 23-25ss zebrafish tailbuds using HCR, and were able to quantify 10 of 13 images (2x 23SS, 3x 24SS and 5x 25SS). As with the cell tracking data, only one PSM was imaged and analysed. We measured nuclear gene expression for genes and signals using an Imaris pipeline, again yielding a set of point clouds describing gene expression and nuclear position for each tailbud. See Materials and Methods for a full description of staining, image acquisition, and quantification.

### AGET construction

AGETs are constructed to approximate the gene expression dynamics of single cells as they move and undergo complex re-arrangements during tissue morphogenesis. This requires live-imaging data, which provides information of the cell’s spatial trajectories over time, to be combined with quantitative single cell gene expression data. To achieve this, we project the pre-processed HCR data onto the tracks to obtain an approximated read-out of the gene expression and signalling levels that each cell experiences as it moves.

We first aligned the point clouds representing the positions of the cells in 3D space processed from the HCRs (Supplementary Figure 1) with the point clouds for each of the 61 time frames in the time lapse (Figure 2A). We use point-to-plane ICP (iterative closest point) to perform this alignment (*30*), which in brief, is an iterative algorithm that seeks to map two point clouds onto each other by recursively minimising the distance between them (see Materials and Methods). Once the point clouds have been aligned, equivalent regions of different PSMs will overlap in space (Figure 2A) making it possible to use the quantitative gene expression from cells in the processed HCRs to assign gene expression values to the cells in the time lapse at each time frame (Figure 2C and D and Algorithm 1 (Materials and Methods)).

**Figure 2:**
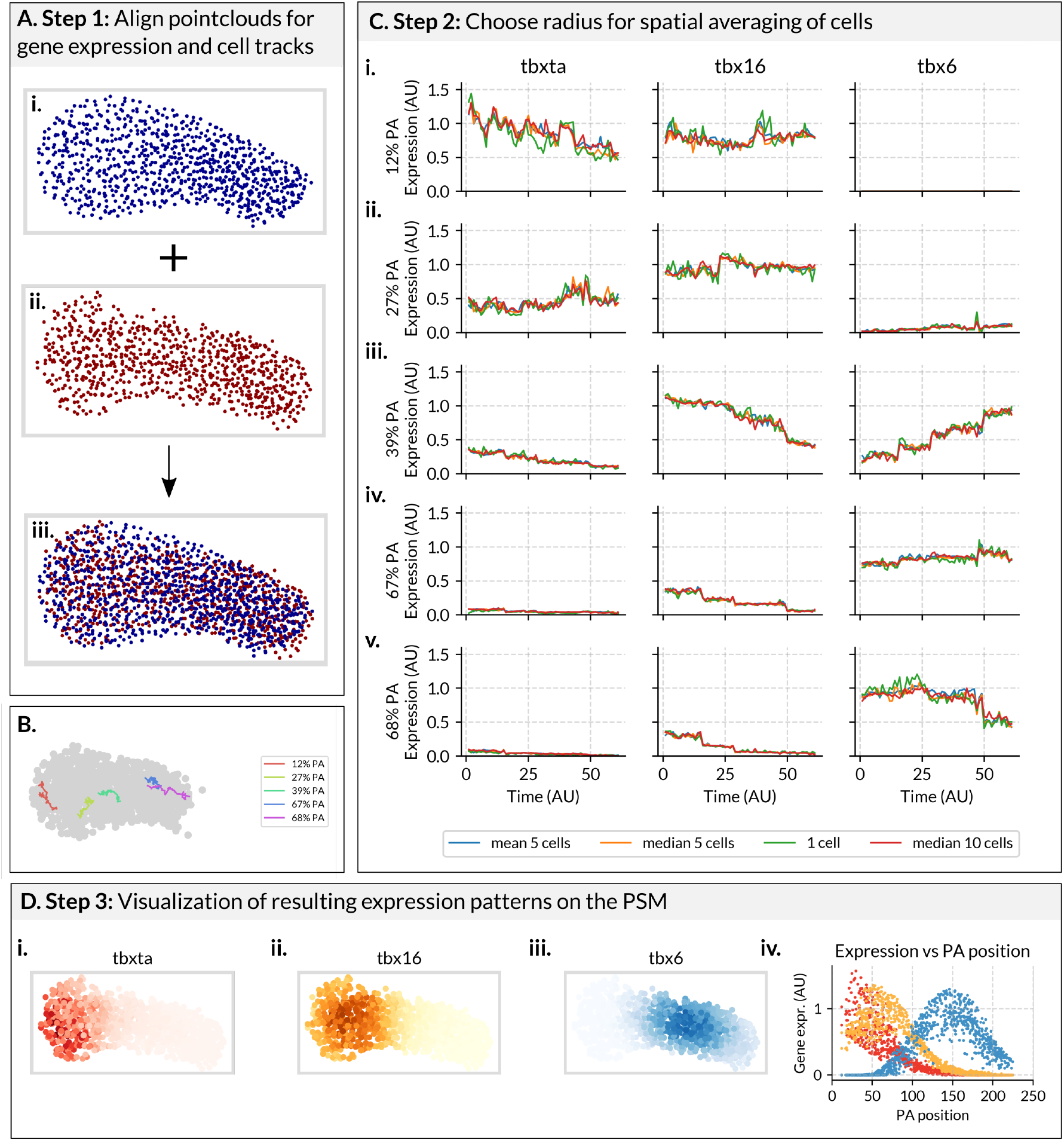
AGET Construction. A. Step 1: Align point clouds representing the positions of cells’ nuclei within the PSM in HCR images and time lapse frames. **A.i**. The point cloud in blue represents the positions of the cells (one point, one cell) in the PSM of a the processed HCR image (source point cloud). **A.ii**. The point cloud in red represents the positions of the cells’ nuclei from the first frame of the tracking data (target point cloud). **A.iii**. Using ICP, all the source point clouds obtained from the HCR images are aligned with the target point cloud obtained from each frame (61 in total) of the tracking data. This is illustrated by the overlapping red and blue point clouds in the resulting point cloud (bottom). **B**. Relative position within the PSM of the AGETs displayed in C. **C. Step 2: Assign a gene expression value to every cell in every frame of the tracks based on the expression of its neighbours from the HCRs**. AGETs representing approximated T-box gene expression dynamics in cells stating at i. 12%, ii. 27%, iii. 39% iv. 67% and v. 68% PA (posterior-to-anterior) position in the PSM. Y-axis represents relative gene expression levels and x-axis, the time frame in the time lapse from 1 to 61. AGETs calculated using different numbers of neighbours and different averaging methods are shown (blue: a cell is assigned the mean expression of its 5 closest neighbours, orange: a cell is assigned the median expression of its 5 closest neighbours, green: a cell is assigned the expression of its closest neighbour, red: a cell is assigned the median expression of its 10 closest neighbours), and from the AGETs it can be seen that there are no major differences. **D. Step 3: Visualise the AGETs on the tracks and compare the resulting tissue-level expression patterns with those measured from the HCRs. D.i**. AGETs for *tbxta* (red) **D.ii**. *tbx16* (yellow) and **D.iii**. *tbx6* (blue) at the first time frame in the movie. **D.iv**. Projection of the the AGETs in D.i-iv. quantified along the posterior to anterior axis of the PSM. Each dot represents the approximated value of a gene (*tbxta* in red, *tbx16* in yellow and *tbx6* in blue) in a cell. Lines represent maximum projection measures of tbox patterns from the HCRs.

To approximate the gene expression and signalling values in a cell from the time lapse, we first find its *n* closest neighbouring cells from the processed HCR data (Figure 2B) and assign it an averaged value based on their expression values. This is repeated for every cell in each of the 61 frames in the time lapse, resulting in an AGET for every cell in the time lapse (Figure 2C). The number *n* of cells that is appropriate to use might be different depending on the specifics of the developing tissue and its cellular architecture. In the case of the zebrafish PSM we found little difference in the resulting AGETs depending on whether these were calculated using 10, 5 or 1 cell-neighbourhoods (Figure 2B and C). The averaging method used (mean or median) had negligible effect in this case (Figure 2B and C). Altogether this suggests that that AGETs calculated for cells in the developing zebrafish PSM are very robust to the methods used to calculate them (Figure 2). The resulting tissue-level patterns were also found to be robust to the number of cells and the averaging method used (Supplementary Figure 2). Once AGETs have been calculated for every cell in the time lapse, we can visualise the resulting pattern at the level of the tissue by plotting the positions of the cells over time and colour-coding them according to their approximated gene expression values (Figure 2D and Supplementary Movies 1 and 2). We refer to these data-sets as *in silico reporters* since they allow us to visualise gene expression dynamics over the course of development (Supplementary Movies 1 and 2). The resulting tissue-level pattern can be quantified and compared to that of the HCRs, revealing a good match between the two (Figure 2D).

### AGETs can be used to reverse-engineer gene regulatory networks that recapitulate pattern formation on a developing tissue

GRN models are often formulated as systems of coupled differential equations where state variables describe the concentrations of the gene products of interest and parameters represent the interactions between genes, as well as other factors such as production and degradation rates. In the case of the T-box genes, there are three state variables representing *tbxta, tbx16* and *tbx6* levels and a total of 24 parameters to be fit (see Materials and Methods). Dynamic data are required to constrain and fit such models, and in this case these will be provided by the AGETs. AGETs will be used as the target expression dynamics for the fitting procedure, where previously directly measured gene expression dynamics would have been used. As with other fitting procedures, an optimal parameter set will be one that minimises the difference between the target and the simulated data. We chose to use a Markov Chain Monte Carlo (MCMC) algorithm (*31*) to use as our parameter sampling method since MCMC has been extensively used and repeatedly validated for GRN inference (*32*). In addition, MCMC and has the advantage of providing a population of candidate networks by approximating the entire posterior distribution for each GRN parameter.

### Optimal fits are obtained using 10% of all AGETs for fitting

We first sought to identify the optimal number of AGETs to fit to: one that is compuationally feasible while producing high quality fits. We fit to various numbers of AGETs, between 10 and 200, and computed the overall quality of the resulting fits for generated parameter sets. We found that while good fits can be obtained when fitting to as few as 10 AGETs, fitting to increased numbers of AGETs increases the proportion of inferred well-fitting parameter sets.

Model fits will improve with the number of AGETs used as reflected by the increase in the mean likelihood (Figure 3A) and in the mean acceptance proportion for each run (Supplementary Figure 4A), decrease in likelihood spread (Figure 3A) and levelling in the mean auto-correlation score for each run as observed when using 200 AGETs for fitting or more (Supplementary Figure 4B). Since the goodness of the fits and the convergence of the MCMC algorithm will increase up to 200 AGETs, plateauing thereafter (Figure 3A and Supplementary Figure 4) we settle on 200 AGETs (approximately 10% of all available AGETs) as the optimal number of AGETs with which to reverse-engineer GRNs for the rest of this study.

**Figure 3:**
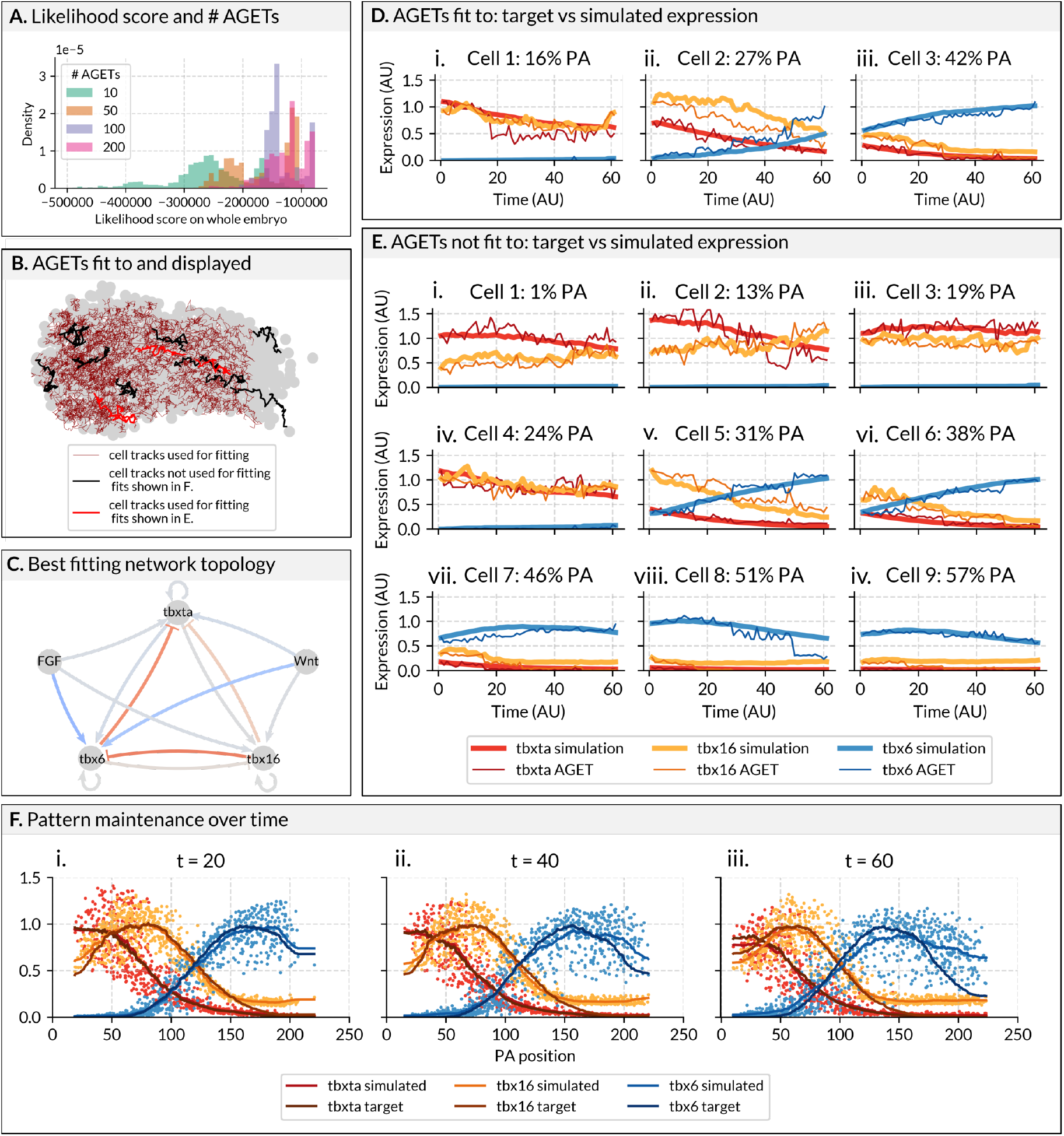
Goodness of fits and performance of the GRN corresponding to the maximum a posteriori (MAP) parameters when fitting to an optimum of 200 AGETs. **A**. Likelihood score improves (increases) with the number of AGETs used for model fitting. GRNs were fit using 10, 50, 100, and 200 AGETs (of a total of around 1903). To compute overall model fit when fitting to different numbers of AGETs, we computed the likelihood using all AGETs for the final 2000 parameter sets of each run (see Materials and Methods). Fitting to 200 cells produced the likelihood distributions with the best mean likelihood values overall, corresponding to the best fitting parameter sets. All GRNs presented from here on were obtained by fitting to 200 AGETs. **B**. Cell tracks whose AGETs have been used for model fitting (200 cell tracks shown in thin red lines), used for model fitting where the fits are shown in D. (3 cell tracks shown in thick red lines) and used for model testing but not fitting, and where the fits are shown in E. (9 cell tracks shown in thick black lines). Cell tracks are shown against the outline of the PSM to convey their relative position within the tissue. **C**. Topology of the best fitting network: this is the parameter set with the lowest likelihood score when computed on all simulated AGETs. Positive interactions are blue arrows, negative interactions are red T-bars, small/zero interaction values are indicated by grey lines. The magnitude of the interaction is indicated by colour intensity. Parameter values are shown in Table 1 (Materials and Methods) **D**. Comparison of simulated vs target AGETs for three cells used in the fitting procedure, which are initially located at i. 12%, ii. 23% and iii. 27% PA within the PSM. Red: *tbxta* expression, yellow: *tbx16* expression, and blue: *tbx6* expression. Thick lines denote simulated gene expression and thin lines, the target AGETs. **E**. Comparison of simulated vs target AGETs for nine cells not used in the fitting procedure, which are initially located at i. 5%, ii. 11%, iii. 12%, iv. 19%, v. 28%, vi. 32%, vii. 36%, viii. 36%, and ix. 47% PA within the PSM. Red: *tbxta* expression, yellow: *tbx16* expression, and blue: *tbx6* expression. Thick lines denote simulated gene expression and thin lines, the target AGETs. **F**. Snapshots showing simulated tbox expression quantified along the PA axis of the PSM at three time frames in the time-lapse: i. 20, ii. 40 and iii. 60. Each dot represents the simulated value of a gene (*tbxta* in red, *tbx16* in yellow and *tbx6* in blue) in a cell at a given time point. Lines represent maximum projection measures of tbox patterns from the HCRs (smoothed average gene expression profiles). Note that the anterior down-regulation of *tbx6* is not recapitulated; this was intentionally not fit to as factors not included in the current GRN formulation are responsible for this feature of the pattern.

**Figure 4:**
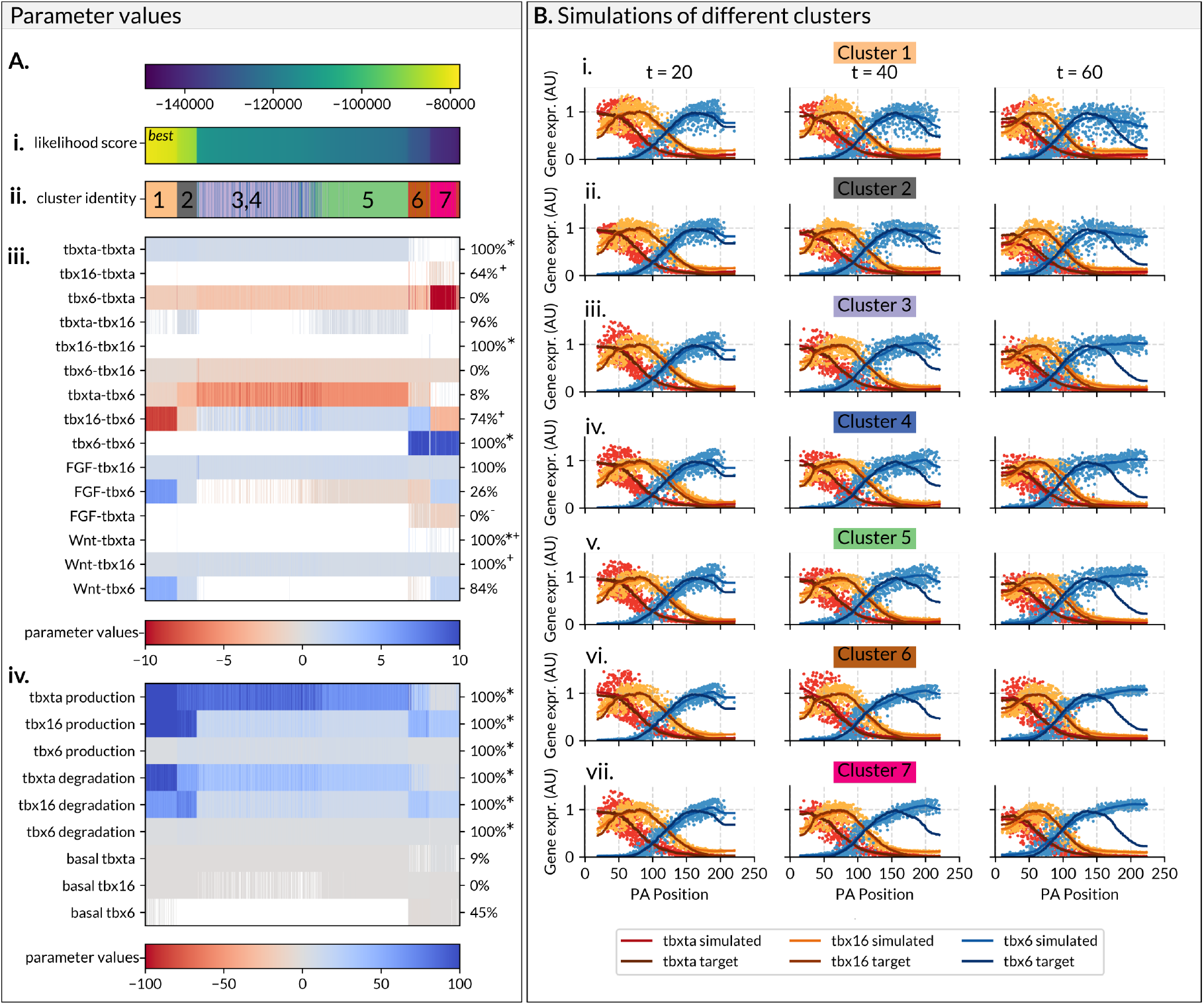
Seven main clusters of GRN topologies that recapitulate Tbox expression along the developing zebrafish PSM. Reverse-engineering GRN topology using sets of 200 AGETs yields seven main clusters of solutions which recapitulate tbox spatio-temporal gene expression along the PSM, as well as known genetic interactions. **A**. The likelihood score for the final converging 2000 sets of parameters is computed on all the tracks. Networks are displayed in order of increasing likelihood (improved fit). **A.i**. Likelihood scores of successful parameter sets. **A.ii**. Cluster identity of parameter sets, from k-means clustering. **A.iii**. Parameters representing interactions between GRN nodes. **A.iv**.. Parameters representing production, degradation and basal gene expression. Parameters indicated by an asterisk (*) are set as positive priors when fitting. **E**. Resulting simulations from representative networks from each cluster.

The 200 AGETs used for model fitting were selected randomly and are distributed uniformly throughout the PSM (Figure 3B, thin red lines). We only selected AGETs from cells that had been consecutively tracked for the entire duration of the time lapse (61 frames). We expect that the optimal number of AGETs required to obtain good fits will be system-specific.

**Table 1:** Parameter values of the MAP network.

|  |  | tbxta | tbx16 | tbx6 |
| --- | --- | --- | --- | --- |
| Inter-connectivity matrix $W$ | tbxta | 0.665 | 0.305 | 1.52 |
|  | tbx16 | -2.27 | 0.00816 | -12.3 |
|  | tbx6 | -10.9 | -0.796 | 0.00110 |
| External Input $E$ | Wnt | 1.66 | 0.511 | 6.63 |
|  | FGF | 0.815 | 0.791 | 8.76 |
| Production rate | $R$ | 28.2 | 160 | 2.73 |
| Degradation rate | $\lambda$ | 5.15 | 87.4 | 1.66 |
| Basal production | $h$ | 0.0612 | -0.923 | -0.899 |

### The MAP network recapitulates Tbox gene expression dynamics at the cellular and tissue levels

MCMC inference yields a collection of parameter sets (samples) that together approximate the posterior distribution of the GRN’s parameters. For every parameter, we obtain a probability distribution across its possible values, which provides information about the values that are most likely to produce good fits. We first chose to explore the network corresponding to the parameter set with the overall highest posterior probability score: the network corresponding to the maximum a posteriori - or MAP - sample (Fig. 3C). We use this network to simulate all 1903 available AGETs, and visualise the simulation on the tracks (Supplementary Movies 2 and 3). We validate the quality of the inferred network by both comparing single AGETs with their simulated counterparts (Fig.3D and E), and by comparing the whole tissue-level gene expression profiles over time (Figure 3F). When simulating single AGETs we find that the MAP network recapitulates both AGETs used for fitting and not used for fitting alike (Figure 3B-D). The model was formulated as a deterministic system without added stochasticity which explains the smoothness of the simulated curves, which nonetheless can be seen to recover AGET gene expression levels and trends.

We are especially interested in how well the simulations recapitulate whole tissue patterning dynamics, as these emerge mostly from simulating AGETs that have not been used for model training (approximately 90% of all AGETs). Figure 3F.i-iii. shows simulated T-box expression for each cell along the normalized posterior to anterior axis of the PSM (dots) at time frames 20, 40 and 60. Simulated data have been fit at each separate time point by curves which are then normalised (darker curves) and compared to the curves previously obtained from the AGETs (shown as lighter curves) (Fig.3F). Overall, simulations recapitulate target tissue-level gene expression very well (snapshots in Fig.3F and full simulations in Supplementary Movies 2 and 3).

Note that there is a discrepancy between the AGETs and the simulated anterior *tbx6* expression. The formulated GRN is unrealistic in this region, as additional factors secreted from the somites are known to be down-regulate this transcription factor (*33*). For this reason we intentionally excluded this region during the parameter optimisation procedure by omitting cells in the anterior-most region of the PSM, and accordingly, simulated gene expression here is inaccurate. In addition, the model predicts that over time, a small percentage of posterior cells will express low levels of *tbx6*. Although unexpected, there is evidence suggesting that this is indeed the case (*26*). Such low and sparse posterior expression of *tbx6* would have been lost during the smoothing step in our data preparation pipeline, which is unable of capturing patterns of such fine resolution as it stands. It is encouraging that candidate GRNs consistently recapitulate this unexpected feature of the biology and might suggest that the three genes considered are indeed causally responsible for much of this patterning system.

### Reverse engineering GRNs using 200 AGETs yields seven clusters of solutions

MCMC is a parameter sampling algorithm, and as such it will return an approximated posterior distribution for the GRN parameters instead of a single point estimate. This provides a range of candidate networks that can be subsequently analysed and challenged in combination with experimental approaches. Such parameter distributions also provide valuable information regarding which model parameters — and therefore genetic interactions — are tightly constrained by the data, and which are not, taking instead a broad range of values across the inferred networks. Such information can lead to interesting hypotheses regarding which aspects of the pattern evolution might be most strongly acting on.

While in the previous section we analysed the network corresponding to the parameter set with the maximal posterior probability (MAP) to asses the goodness of fit of one of the candidate GRNs, in this section we assess how well the posterior distribution has been approximated across candidate GRNs (Figure 5). To do this, we calculated the likelihood score across all the tracks (Materials and Methods) for the final 2000 parameter sets for every fitting run which reached convergence. As the MCMC algorithm used can produce small numbers of poorly fitting parameter sets, even when convergence has been reached, we then excluded all parameter sets with an overall likelihood score of less than -15000, as well as parameter sets with unrealistic values (genetic interactions *<* 100 or *> −*100). This left us with a total of 3600 candidate networks. We plotted these networks in order of increasing likelihood (better fit) (Figure 5A-D) colour coding according to whether a give interaction is positive (activation, red) or negative (repression, blue) to visualise the different predicted network topologies.

**Figure 5:**
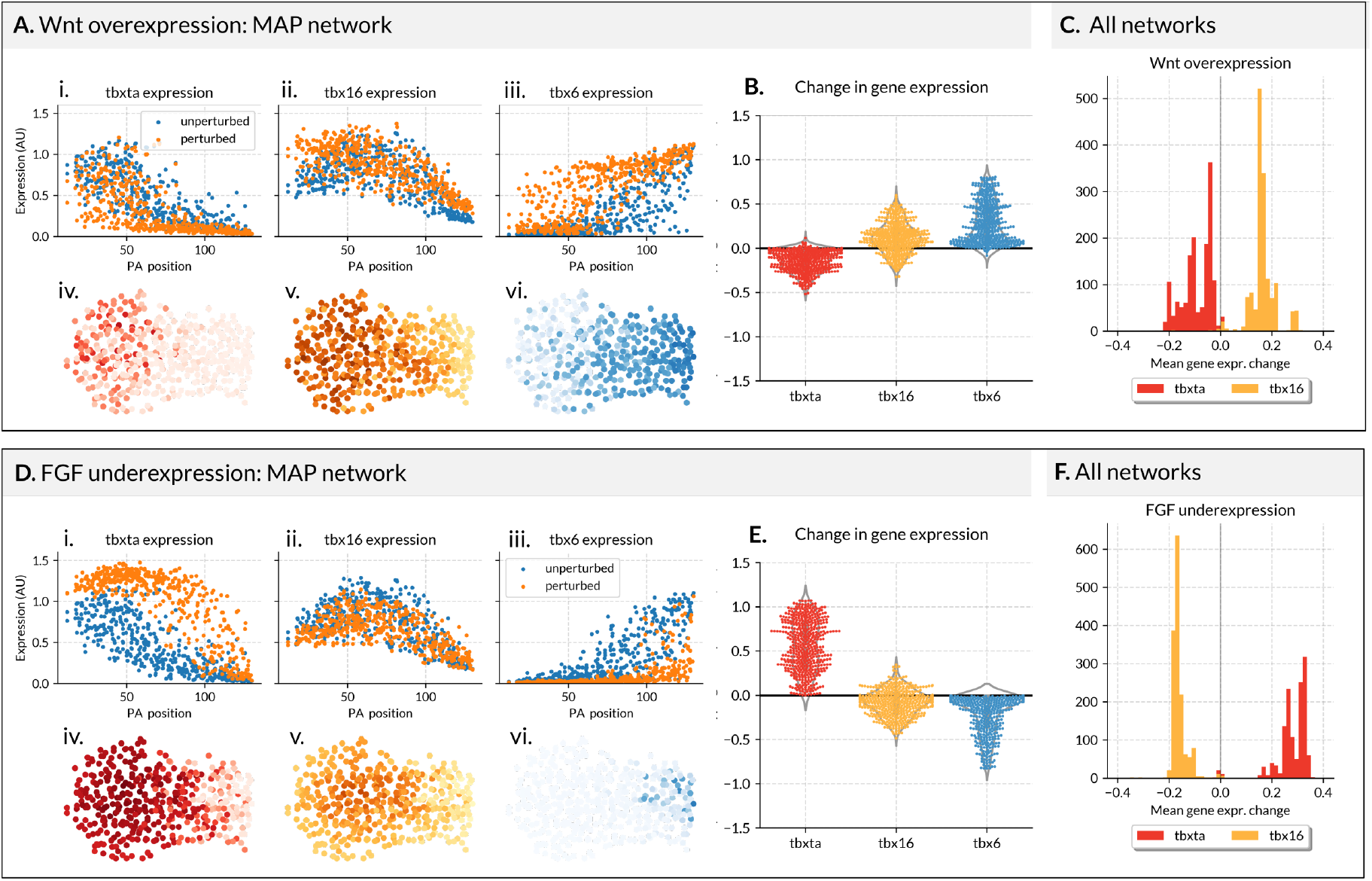
GRNs reverse-engineered using 200 AGETs recapitulate known results of experimental perturbations. **A**. Results of simulating Wnt over-expression using the MAP network. **A.i**. *tbxta*, **A.ii**. *tbx16* and **A.iii**. *tbx6* expression in unperturbed (blue) and perturbed (orange) simulations using the MAP network. X-axis denotes posterior to anterior position along the PSM. Y-axis represents gene expression levels in arbitrary units. Each dot represents the expression level in a cell at the last time point of the simulation. **A.iv**. *tbxta* (red), **A.v**. *tbx16* (yellow) and **A.vi**. *tbx6* (blue) expression shown in cells within the PSM. Posterior left, dorsal up. **B**. Difference in gene expression between the unperturbed and perturbed simulations calculated for every cell at the last time point of the simulations. **C**. The mean gene expression difference in single cells between perturbed and unperturbed simulations, calculated for every network and displayed as histogram for *tbxta* (red) and *tbx16* (yellow). **F**. Results of simulating FGF under-expression. **D.i**. *tbxta*, **D.ii**. *tbx16* and **D.iii**. *tbx6* expression in unperturbed (blue) and perturbed (orange) simulations using the MAP network. X-axis denotes posterior to anterior position along the PSM. Y-axis represents gene expression levels in arbitrary units. Each dot represents the expression level in a cell at the last time point of the simulation. **D.iv**. *tbxta* (red), **D.v**. *tbx16* (yellow) and **D.vi**. *tbx6* (blue) expression shown in cells within the PSM. Posterior left, dorsal up. **E**. Difference in gene expression between the unperturbed and perturbed simulations calculated for every cell at the last time point of the simulations. **F**. The mean gene expression difference in single cells between perturbed and unperturbed simulations, calculated for every network and displayed as histogram for *tbxta* (red) and *tbx16* (yellow).

We clustered these networks using k-means clustering on scaled parameter values (Materials and Methods). We initially clustered them into 13 clusters in total (as determined by an elbow plot - supplemental data), and considered only those clusters which included 5% or more of the total number of networks, reducing the number of clusters to seven. All clusters produce well-fitting patterns (Figure 5B). Furthermore, since all seven clusters of networks have very similar likelihood scores, we treat them all as equally probable candidates until further biological insight helps us to discriminate between them (*26*). We are confident that our methodology is effective at producing a range of topologically distinct and well fitting parameter sets, as evidenced by the accuracy of the resulting fits for all of the networks.

We investigated whether these networks recovered genetic interactions known from the literature. For instance, Wnt is known to directly activate *tbx16* in the zebrafish tailbud: Bouldin et al. identify a Wnt-binding promoter that drives *tbx16* expression (*34*)/ All our parameter sets predict this interaction is positive, thus being supported by the literature (Figure 5A.iii.). *tbx16* is predicted to activate *tbx6* in 74% of the inferred networks (Figure 5A.iii.). The nature of this interaction has been experimentally demonstrated using heat-shock transgenic lines to up-regulate *tbx16* expression which led to an increased expression of *tbx6* (*34*). *tbx16* mutants have a loss of *tbx6* expression (*35*), further corroborating this interaction. (*34*) also show that over-expression of *tbx16* results in a decrease of *tbxta* in the tailbud, but that this phenotype is rescued by over-expression of Wnt, suggesting that this is an indirect interaction occurring via Wnt. 64% of our parameter sets have a positive interaction between *tbx16* and *tbx6* (Figure 5A.iii.). Similarly, (*36*) show that at the tailbud stage, loss of FGF causes an expansion of *tbxta*, suggesting that FGF inhibits *tbxta*; in all of our parameter sets, FGF inhibits *tbxta*. It is also known that FGF activates *tbx16* expression as dominant negative FGFR1 embryos display a loss of *tbx16* expression (***?***, *36*); again, is all of our parameter sets FGF is activating *tbx16* (Figure 5A.iii.). Together, these data reveal a high degree of consensus between our parameter values and known interactions established in the literature.

### Reverse-engineered GRNs qualitatively recapitulate known results of experimental perturbations

In the previous section we validated our parameter sets and GRN inference methodology by comparing our parameter sets to known genetic interactions from the literature. To further validate our parameter sets, we attempted to replicate the effects of changing FGF and Wnt signalling. Bouldin et al. over-express Wnt in the zebrafish tailbud by over-expressing beta-catenin under a heat-shock promoter. Four hours post heat-shock, the *tbx16* expression domain is expanded towards the anterior PSM as a result of an up-regulation of *tbx16* in this region. We replicated this experiment in silico by setting Wnt expression to 1.5 and maintained it for the duration of the simulation. We then ran these simulations on the tracks as before. When this simulation was run using the MAP network we find that at the final time-point of simulation *tbx16* is up-regulated in the anterior PSM, as in the experimental data (Fig. 1A and B. We see, however, a down-regulation of *tbxta* which is opposite to the up-regulation of *tbxta* reported by Bouldin et al. This could be related to the fact that our model formulation does not include the known feedback loop between Wnt and *tbxta* (*37*). When we run this in silico experiment with all the networks we find that the vast majority predict an average increase in *tbx16* expression (Figure 1C).

Goto et al. downregulate FGF expression by inducing a heat-shock dominant negative form of FGF at 8 ss (*36*). At this time point, loss of FGF results in an expansion of *tbxta* expression three hours after heatshock, at approximately the 15ss. We reproduced this experiment by setting FGF to 0.01 in every cell and simulating as before. For the MAP network, *tbxta* is significantly up-regulated after downregulation of FGF, in agreement with the experimental results. *tbx16* is slightly down-regulated, and *tbx6* is significantly down-regulated (Figure 1DE). When we repeated this analysis on all successful parameter sets, a majority recapitulate an increase in *tbxta* expression, as represented by the average difference between *tbxta* expression in each cell in WT and perturbed simulations (Figure 1F).

## Conclusion

Earlier reverse-engineering frameworks have been unable to accommodate the role of cell rearrangements and tissue shape changes in developmental pattern formation. This limitation has heavily biased quantitative studies of pattern formation towards systems where the timing of pattern formation and morphogenesis can be separated. However, the vast majority of patterning processes in animal development do not meet this criterion and in consequence, their study has been grossly under-represented in the GRN literature. As a result, most of our collective knowledge and understanding of the generation and evolution of developmental patterns has been constructed on the omission of any role that might be played by cell movements, tissue shape changes and other morphogenetic mechanisms.

Here, we propose a method to reverse-engineer gene regulatory networks using datasets that are relatively straightforward to generate in an increasing number of model and non-model species spanning the range of animal phylogeny. This will make it possible to construct AGETs and therefore infer GRNs in a wider range of systems. Simulation and subsequent analysis of patterning processes that are dependent on or, at least, co-occurring with cell movements will increase our understanding of pattern formation and its evolution, and uncover general principles that were inaccessible with previous approaches. Furthermore, this methodology will find applications well-beyond beyond the study of developmental evolution. In particular, we anticipate technique will be particularly useful in fields such as bio-engineering, regenerative medicine and organoid biology, where understanding how 3D cell cultures should be shaped and constrained as they grow to obtain the desired final organisation is paramount and has proven not at all trivial.

While our methodology is in principle applicable across a wide variety of developmental processes, it would have to be substantially modified to accommodate highly dynamic patterning processes. Our methodology makes the assumption that a cell’s gene expression is strongly correlated with its position within a tissue. This is an assumption that applies well to Tbox patterning in the zebrafish PSM, where cells are known to express the gene sequence *tbxta, tbx16, tbx6* as they differentiate and transit towards the somites (*26*). However, this assumption does not always apply; for example oscillatory gene expression in during somitogenesis (*38*) or neural differentiation (*39*). In such systems, our current AGET construction methods would need to be revised and HCRs would need to be cover the dynamics of of the process at a much higher resolutioin. Still, since this method is based on approximating dynamics from static images, one would risk missing very rapid or transient gene expression dynamics. To mitigate this, we have performed extensive model validation using features of the GRN that have not been fit to, such as GRN topology, and functional analysis/ability to respond to perturbations.

Finally, our methodology for the construction of AGETs provides a way in which to visualise approximated gene expression dynamics and patterning in the form of in-silico reporters. There is in principle no limit to the number of genes that can be reported by an in silico reporter line which could be used for hypothesis generation and to compare the relative co-expression of previously unexplored combinations of genes. In silico reporter lines can also be readily extended to non-model organisms where transgenic reporters are not yet available. All in all, we expect that this methodology will find a broad range of applications in developmental evolution and beyond, contributing to advance the data-driven study of patterning dynamics.

## Materials and Methods

### Animal lines and husbandry

This research was regulated under the Animals (Scientific Procedures) Act 1986 Amendment Regulations 2012 following ethical review by the University of Cambridge Animal Welfare and Ethical Review Body (AWERB). Embryos were obtained and raised in standard E3 media at 28°C. Wild Type lines are either Tupfel Long Fin (TL), AB or AB/TL. The Tg(7xTCF-Xla.Sia:GFP) reporter line (*40*) was provided by the Steven Wilson laboratory. Embryos were staged as in (*41*).

### In Situ Hybridisation Chain Reaction (HCR)

Embryos were incubated until they reached the the desired developmental stage, then fixed in 4% PFA in DEPC treated PBS without calcium and magnesium, and stored at 4°C overnight. Once fixed, embryos were stained using HCR version 3 following the standard zebrafish protocol found in (*29*). Probes, fluorescent hairpins and buffers were all purchased from Molecular Instruments. After staining, samples were stained with DAPI and mounted using 80% glycerol.

### Immunohistochemistry

Embryos were incubated until they reached the desired developmental stage, then fixed in 4% PFA in DEPC treated PBS without calcium and magnesium, and stored at 4°C overnight. The embryos were subsequently blocked in 3% goat serum in 0.25% Triton, 1% DMSO, in PBS for one hour at room temperature. Our read-out for FGF activity - Diphosphorylated ERK - was detected using the primary antibody (M9692-200UL, Sigma) diluted 1 in 500 in 3% goat serum in 0.25% Triton, 1% DMSO, in PBS. All samples were incubated at 4°C overnight and then washed in 0.25% Triton, 1% DMSO, in PBS. Secondary Alexa 647nm conjugated antibodies were diluted 1 in 500 in 3% goat serum in 0.25% Triton, 1% DMSO, 1X DAPI in PBS and applied overnight at 4°C.

### Imaging and image analysis

Fixed HCR and immunostained samples were imaged with a Zeiss LSM700 inverted confocal at 12 bit, using either 20X or 40X magnification and an image resolution of 512x512 pixels. Nuclear segmentation of whole stained embryonic tailbuds was performed using a tight mask applied around the DAPI stain using Imaris (Bitplane) with a surface detail of 0.5µm. Positional values for each nucleus were exported as X, Y, Z coordinates relative to the posterior-most tip of the PSM where X, Y, Z were equal to (0, 0, 0). The PSM was then segmented by hand by deleting nuclear surfaces outside of the PSM, including notochord, spinal cord, anterior somites and ectoderm. PSM length was normalised individually between 0 and 1 by division of the position in X by the maximum X value measured in each embryo.

Single cell image analysis was conducted using Imaris (Bitplane) by generating loose surface masks around the DAPI stain to capture the full nuclear region and a small region of cytoplasm. Surface masks were then filtered to remove any masks where two cells joined together or small surfaces caused by background noise, or fragmented apoptotic nuclei. The intensity sum of each channel was measured and normalised by the area of the surface. Expression level was then normalised between 0 and 1 using the maximum value measured for each gene, in each experiment.

Live imaging datasets of the developing PSM were created using a TriM Scope II Upright 2-photon scanning fluorescence microscope equipped Insight DeepSee dual-line laser (tunable 710-1300 nm fixed 1040 nm line) (see details in (*27*)). The developing embryo was imaged with a 25X 1.05 NA water dipping objective. Embryos were positioned laterally in low melting agarose with the entire tail cut free to allow for normal development (*42*). Tracks were generated automatically and validated manually using the Imaris imaging software.

### Image analysis processing pipeline for HCR images

The first step in the pipeline consists of masking the PSM from the surrounding tissues, including the spinal cord and the notochord. This was achieved by drawing a surface around the PSM using morphological and gene expression landmarks as a guide to identify different tissue boundaries (Supplementary Figure 1B). Next, in order to consider only gene expression levels inside of the isolated PSM, all gene expression outside of the defined surface was set to zero (Supplementary Figure 1C). Background noise in the data was reduced by setting lower-bound thresholds for every gene. These thresholds were chosen such that *tbxta* and *tbx16* would appear restricted to the posterior end of the PSM (Supplementary Figure 1Di, Dii, Ei and Eii) with their expression in the anterior PSM reduced to zero. Similarly, thresholds were set for *tbx6* expression to eliminate any background expression in the posterior PSM (Supplementary Figure 1Diii and Eiii). Each gene is then normalized; normalization had to be robust enough to noisy gene expression levels. A Savitzky-Golay filter was applied to each gene to smoothen the signal (Fig.1D) and the smoothened maximum for each gene was set to one. Finally, spots were created in each detected nucleus from which a point cloud consisting of the 3D spatial coordinates and associated *tbxta, tbx16* and *tbx6* levels were extracted (Supplementary Figure 1E). The same pipeline was used to obtain the levels of signals Wnt and FGF in single cells.

### Aligning point clouds with ICP

We used the Python library Open3d (*43*) and the implementation of the point-to-plane ICP (Iterative Closest Point) algorithm therein (*30*) to perform the point cloud alignment. ICP algorithms can be used to align two point clouds from an initial approximate alignment. The aim is to find a transformation matrix, that rotates and moves the source point cloud in a way that achieves an optimal alignment with the target point cloud. ICP algorithms work by iterating two steps. First, for each point in the source point cloud, the algorithm will determine the corresponding closest point in the target point cloud. Second, the algorithm will find the transformation matrix that most optimally minimizes the distances between the corresponding points. The result is a transformed source point cloud that is closely aligned with the target point cloud. As a pre-processing step, the source and target point clouds have been re-scaled to have the same A-P length. Since we are working with biological tissues, point clouds will not correspond exactly, differing slightly in size and shape. This will impact the quality of the resulting alignment which had to be visually assessed and validated. In this case study, three of the thirteen source images were excluded from the analysis due to poor alignment.

### AGET construction

While the main methodology used for constructing AGETs is covered in the results section, below (Algorithm 1) we provide pseudo-code that describes the same process.

#### Algorithm 1

Mapping T-box gene expression from HCR images onto tracking data

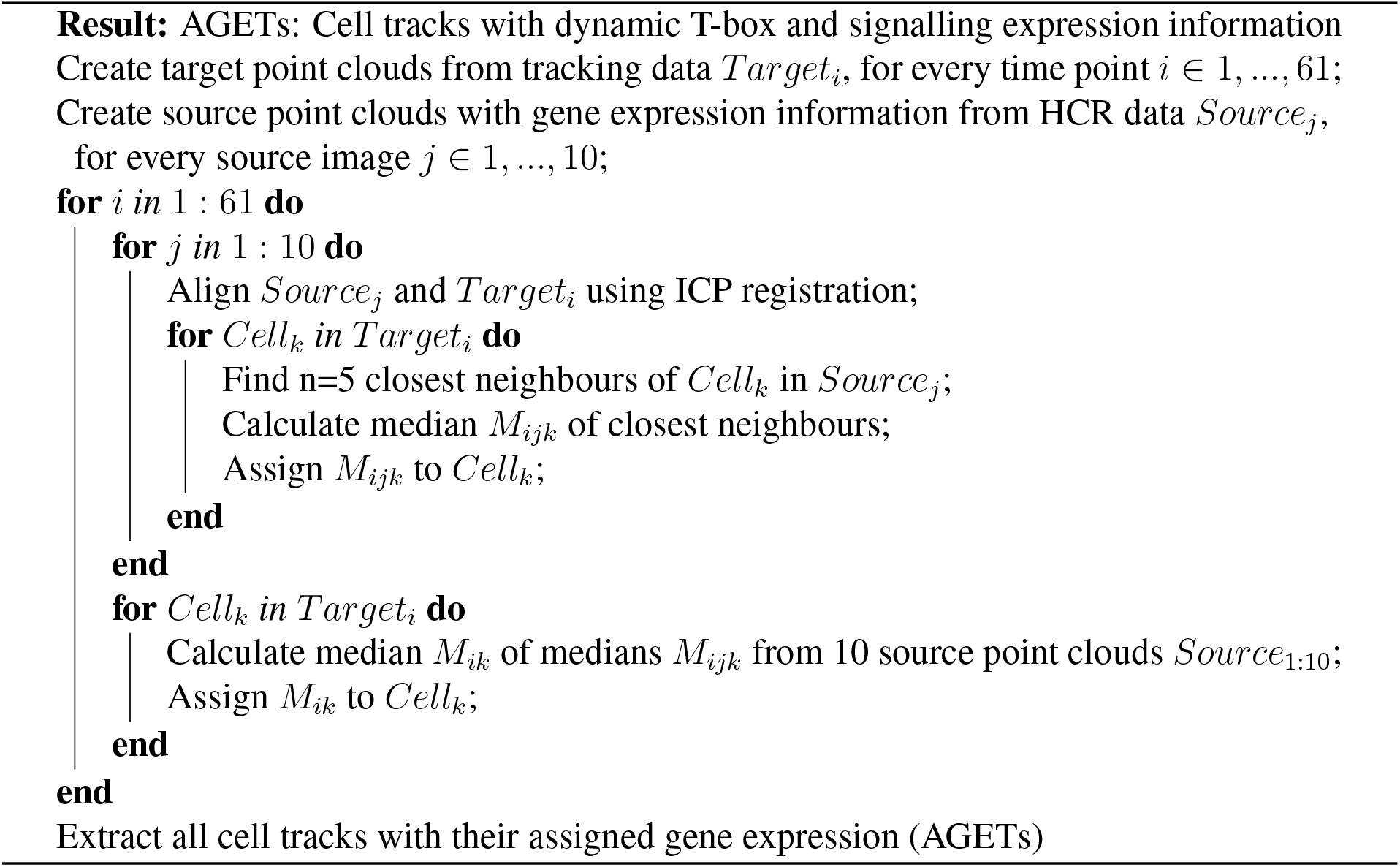

### Mathematical model formulation

We used a dynamical systems formulation model the T-box gene regulatory network in the zebrafish PSM. The model’s aim is to recapitulate the dynamics of T-box gene expression in every cell in the developing zebrafish PSM, generating the emergence of the tissue-level T-box gene expression pattern. We use a connectionist model formulation which has been extensively used and validated to previously model other developmental patterning processes (*8, 14, 44*).

The mRNA concentrations encoded by the T-box genes *tbxta, tbx16* and *tbx6* are represented by the state variables of the dynamical system. For each gene, the concentration of its associated mRNA *a* at time *t* is given by *g*^*a*^(*t*). mRNA concentration over time is governed by the following system of three coupled ordinary differential equations:

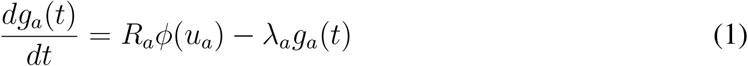

where *R*_*a*_ and *λ*_*a*_ respectively represent the rates of mRNA production and decay. *ϕ* is a sigmoid regulation-expression function used to represent the cooperative, saturating, coarse-grained kinetics of transcriptional regulation and introduces non-linearities into the model that enable it to exhibit complex dynamics:

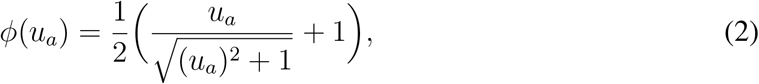

where

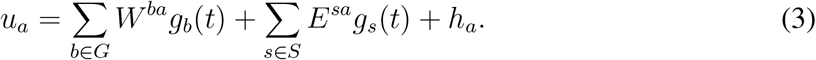

*G* = *{tbxta, tbx16, tbx6}* refers to the set of T-box genes while *S* = {Wnt, FGF} represents the set of external regulatory inputs provided by the Wnt and FGF signalling environments. The concentrations of the external regulators *g*_*s*_ are provided directly from the AGETs into the simulation and are not themselves being modelled. Changing Wnt and FGF concentrations over time renders the parameter term 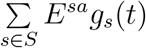 time-dependent and therefore, the model non-autonomous (*45, 46*).

The inter-connectivity matrices *W* and *E* house the parameters representing the regulatory interactions among the T-box genes, and from Wnt and FGF to the T-box genes, respectively. Matrix elements *w*^*ba*^ and *e*^*sa*^ are the parameters representing the effect of regulator *b* or *s* on target gene *a*. These can be positive (representing an activation from *b* or *s* onto *a*), negative (representing a repression), or close to zero (no interaction). *h*_*a*_ is a threshold parameter denoting the basal activity of gene *a*, which acknowledges the possible presence of regulators absent from our model. To perform the live-modelling simulations, the same model formulation is implemented in each cell in the time-lapse. Initial concentrations of *tbxta, tbx16* and *tbx6* are read out directly from the first time point of the AGET corresponding to that cell, and dynamic Wnt and FGF values are updated from the same AGET.

### Model fitting: MCMC approach

We used the Markov Chain Monte Carlo approach implemented in the Python emcee library (*31*) to approximate the posterior distribution of the GRN parameters. A property of this implementation is the use of an ensemble of walkers, rather than a single one. To fit, we used a uniform prior from -200 to +200, except when the priors were further restricted to positive values only, and fitted to the time scale used in the simulation. The time scale was chosen such that 1 equals the time that the fastest cell takes to travel through the whole PSM and enter a somite. We used a Gaussian distribution with fixed standard deviations per gene to model the differences between simulated gene expression and target gene expression approximated by the AGETs, and in this way obtain a likelihood function. We ran the MCMC with 96 walkers and for a total of 10’000 steps. We ran the MCMC independently at least 5 times for each parameter set, as this sometimes obtained different parameter distributions. Each fitting run was then extended for a further 10’000 iterations if it had not reached convergence. We defined convergence using the K-S test to check that the parameter distributions from the final 5000 parameter sets were statistically distinguishable from the -10000th to -8000th parameter sets, for at least 20 of the parameters. We used a significance threshold of 0.001.

### Calculating a likelihood score

To compute the log-likelihood score during MCMC, we computed the squared distance between the target (AGET) and simulated gene expression for all cells fit to. This value was normalized using a custom *σ* value for each gene. More specifically, we used the following likelihood formulation:

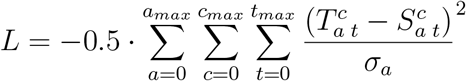

Here, *a* refers to the gene being fit to, *c* refers to the cell being fit to; in this study *c*_*max*_ was between 10 and 200. *t* refers to the time point of the simulation; in this study *t*_*max*_ = 61. *T* refers to the Target - AGET - gene expression, and *S* refers to simulated gene expression. *σ* refers to the standard deviation which was unique for each gene. *σ*_*tbxta*_ = 0.2, *σ*_*tbx*16_ = 0.2, *σ*_*tbx*6_ = 0.1.

### Filtering and clustering

We filtered parameter sets for further analysis as follows. We firstly only analysed parameter sets from runs that had reached convergence, and were fit to 200 AGETs. We then excluded unrealistically large parameter values, keeping values between +-100. We finally excluded parameter sets with a likelihood score less than -15000. Unfiltered parameter sets are presented in Supplementary Figure 7 and filtered in Supplementary Figure 6.

We performed k-means clustering of parameter values filtered as above, using the ‘Kmeans’ function of the ‘sklearn’ library v0.24.2. We chose the number of clusters to use using an Elbow Plot (FIG). We then removed extremely small clusters (less than 5% of all parameter sets) to obtain the seven clusters of parameter values presented in this paper.

## Acknowledgments

The authors would like to thank the Cambridge Advanced Imaging Centre (CAIC) for imaging support and the University of Oxford Advanced Research Computing (ARC) facility for their computational support.

## Contributions

Conceptualisation: BP, BS and BV. Methodology: KS, BV, BS and BP. Software: KS, SET, KT, DS, SH, YW, BP and BV. Validation: KS and SET. Formal Analaysis: KS and SET. Investigation (experimental work): TF. Writing-original draft preparation: KS and BV. Writing-review and editing: BS and BV. Supervision: BP, BS and BV. Funding acquisition: BP, BS and BV.

## Data and Code Availability

Data and code available from https://github.com/spikay/AGETS

---

**Supplementary Fig. 1:**
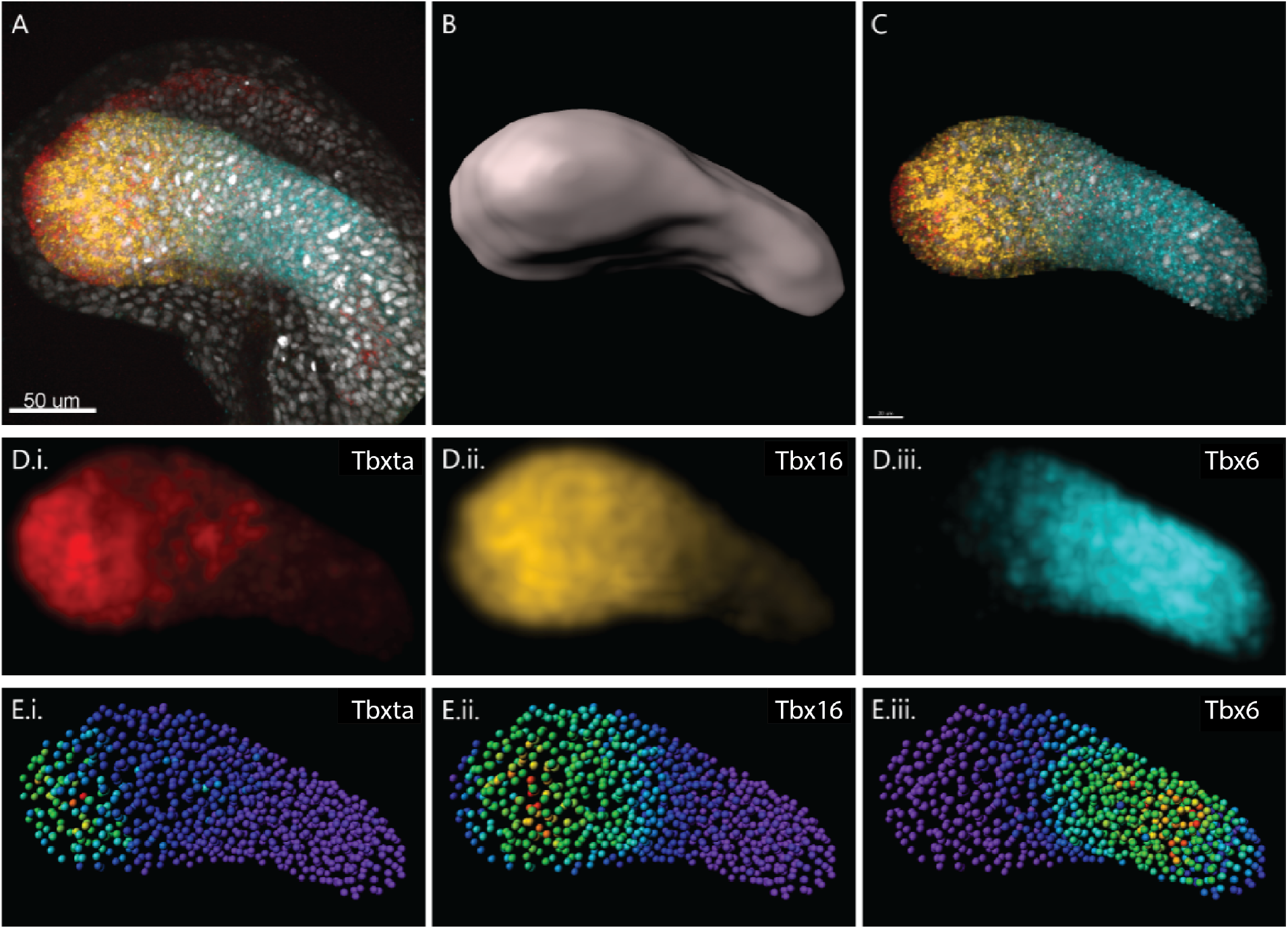
Gene expression data preparation pipeline. **A**. Typical HCR image of a 22 somite stage zebrafish embryo tailbud stained for *tbxta* (red), *tbx16* (yellow), *tbx6* (blue) and DAPI (gray). Anterior to the right, posterior to the left, dorsal up and ventral down from here on. **B**. Surface masking the PSM based on T-box gene expression and morphological landmarks. **C**. Gene expression and nuclear marker in the masked PSM (as before *tbxta* in red, *tbx16* in yellow, *tbx6* in blue and DAPI in gray). **D**. Normalising gene expression levels: *tbxta* and *tbx16* levels in the anterior PSM are normalised to zero while posterior PSM levels of *tbx6* are normalised to zero, to eliminate background expression. A Gaussian filter has been then applied to each T-box gene to smoothen gene expression across the PSM. **E**. Nuclei are segmented using the DAPI channel creating spots in 3D space. Spots are coloured according to the median intensity of each gene i. *tbxta*, ii. *tbx16* and iii. *tbx6*), where purple denotes zero expression and red 1, which is the highest expression. The spatial coordinates of the spots together with the median intensities were exported and used to generate the AGETs.

**Supplementary Fig. 2:**
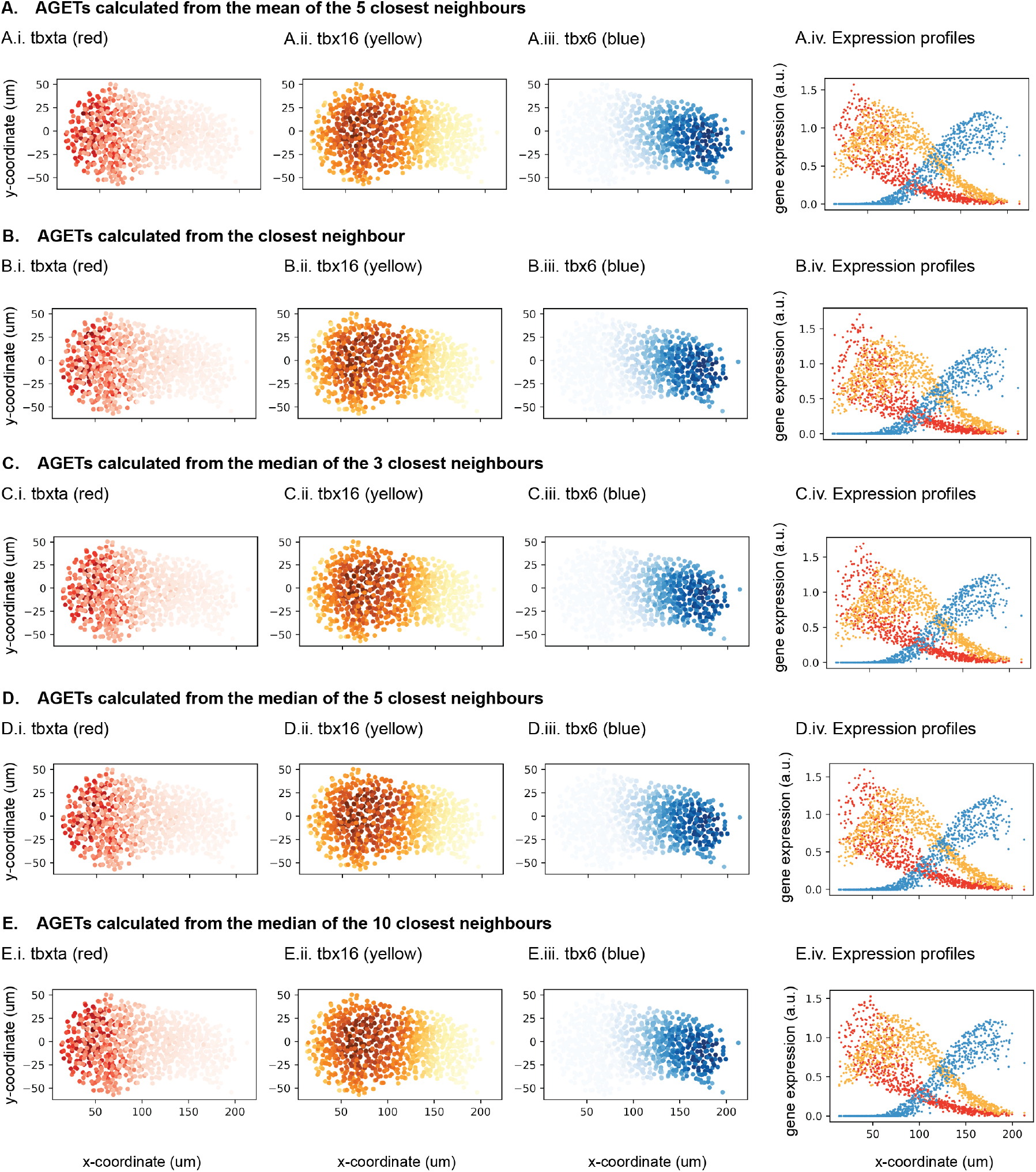
Tissue-level pattern is robust to AGET calculation method. **A**. Approximated Tbox gene expression pattern on the PSM when AGETs were calculated taking the mean of the five nearest neighbours. **A.i**. Approximated *tbxta* in the cells of the PSM. Each dot represents the position of a cell in the PSM (x,y-projection where dorsal is to the top and posterior is to the left). Shade of red indicates approximated *tbxta* concentration (dark red, highest, white, lowest). **A.ii**. Approximated *tbx16* in the cells of the PSM. Shade of yellow indicates approximated *tbx16* concentration (dark yellow, highest, white, lowest). **A.iii**. Approximated *tbx6* in the cells of the PSM. Shade of blue indicates approximated *tbx16* concentration (dark blue, highest, white, lowest). **A.iv**. Tbox gene expression profiles. Each dot represents the concentration of one of the tbox genes (*tbxta* (red), *tbx16* (yellow) and *tbx6* (blue) in a given cell. The position along the posterior to anterior axis of each cell is given by its x-coordinate. **B**. Approximated Tbox gene expression pattern on the PSM when AGETs were calculated taking the value of the nearest neighbour.**B.i**. Approximated *tbxta* in the cells of the PSM. **B.ii**. Approximated *tbx16* in the cells of the PSM. **B.iii**. Approximated *tbx6* in the cells of the PSM. **B.iv**. Tbox gene expression profiles. **C**. Approximated Tbox gene expression pattern on the PSM when AGETs were calculated taking the value of the nearest neighbour.**C.i**. Approximated *tbxta* in the cells of the PSM. **C.ii**. Approximated *tbx16* in the cells of the PSM. **C.iii**. Approximated *tbx6* in the cells of the PSM. **C.iv**. Tbox gene expression profiles. **D**. Approximated Tbox gene expression pattern on the PSM when AGETs were calculated taking the value of the nearest neighbour.**D.i**. Approximated *tbxta* in the cells of the PSM. **D.ii**. Approximated *tbx16* in the cells of the PSM. **D.iii**. Approximated *tbx6* in the cells of the PSM. **D.iv**. Tbox gene expression profiles. **E**. Approximated Tbox gene expression pattern on the PSM when AGETs were calculated taking the value of the nearest neighbour.**E.i**. Approximated *tbxta* in the cells of the PSM. **E.ii**. Approximated *tbx16* in the cells of the PSM. **E.iii**. Approximated *tbx6* in the cells of the PSM. **E.iv**. Tbox gene expression profiles.

**Supplementary Fig. 3:**
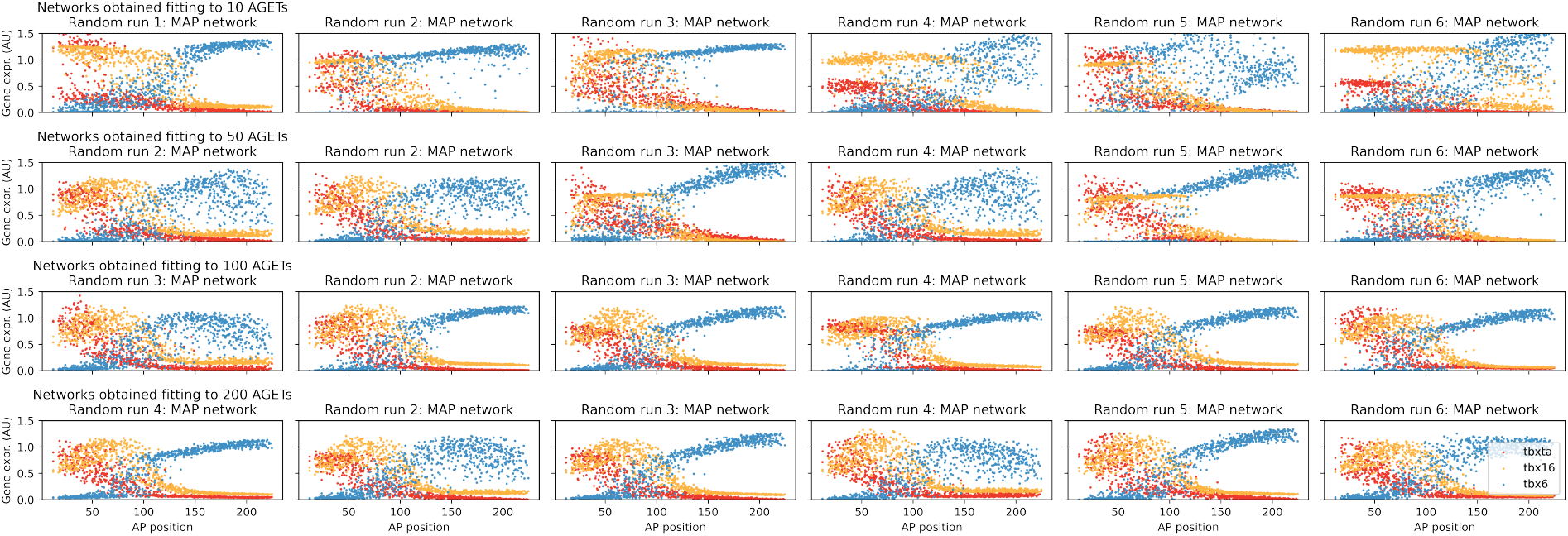
The proportion of parameter combinations producing good fits increases as the number of AGETs used for fitting is increased. **(A)** Networks obtained fitting to 10 AGETs. **(B)** Networks obtained fitting to 50 AGETs. **(C)** Networks obtained fitting to 100 AGETs. **(D)** Networks obtained fitting to 200 AGETs. In each case, the MAP parameter set is taken from 6 independent random runs and the expression profile corresponding to the last time point in the simulation is plotted. Each dot represents the concentration of one of the tbox genes (*tbxta* (red), *tbx16* (yellow) and *tbx6* (blue) in a given cell. The position along the posterior to anterior axis of each cell is given by its x-coordinate. Acceptable fits are obtained regardless of the number of AGETs used for fitting, but the proportion of acceptable fits increases with the number of AGETs.

**Supplementary Fig. 4:**
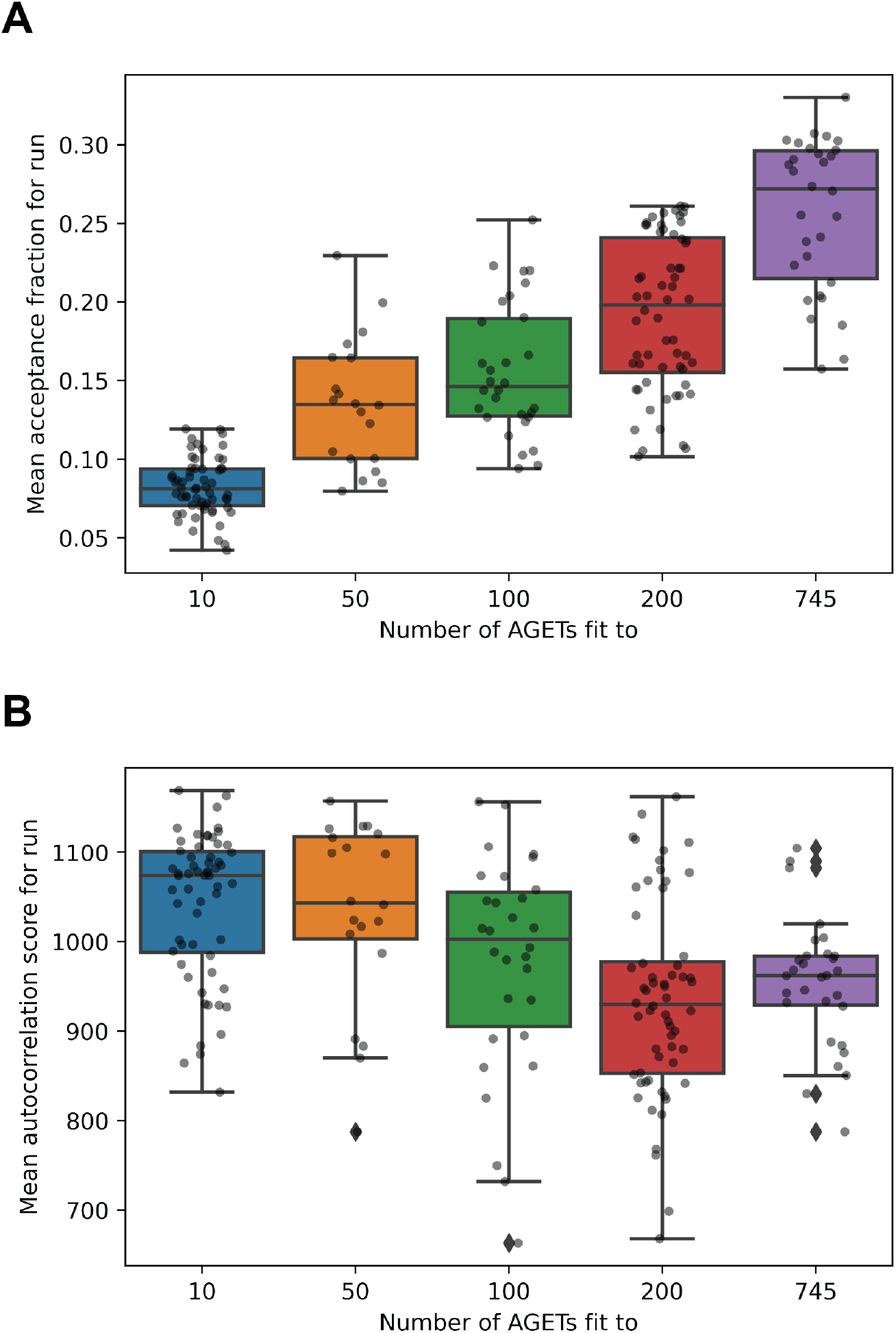
**A**. Mean acceptance fraction per run increases with the number of AGETs used for fitting in initial tests. **B**. Mean auto-correlation score per run decreases as the number of AGETs increases until 200 AGETs, and stabilises thereafter.

**Supplementary Fig. 5:**
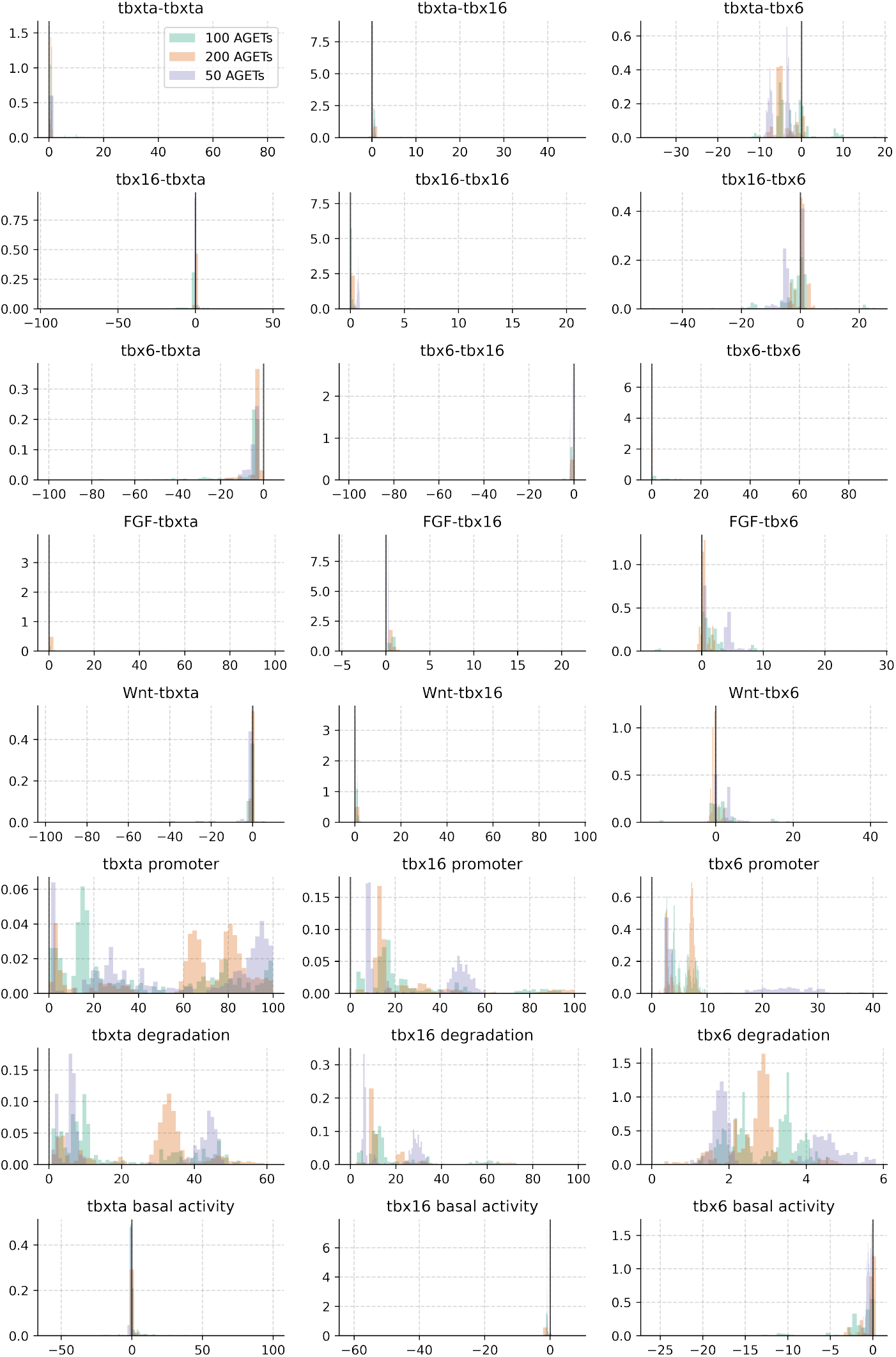
Marginal posterior distribution parameter values obtained using 10, 50, 100 and 200 AGETs.

**Supplementary Fig. 6:**
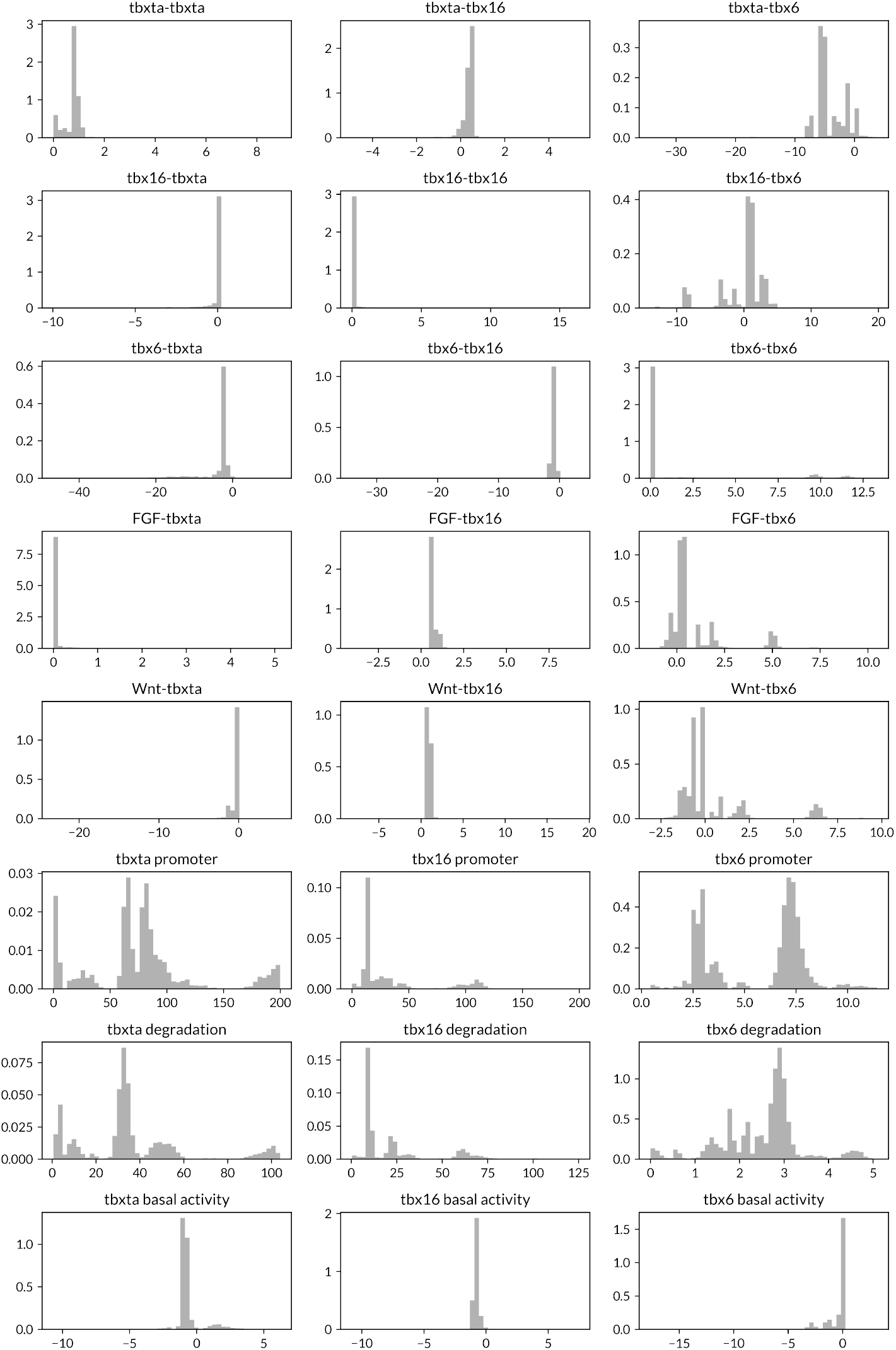
Marginal posterior distribution of filtered parameter values used for analysis and clustering.

**Supplementary Fig. 7:**
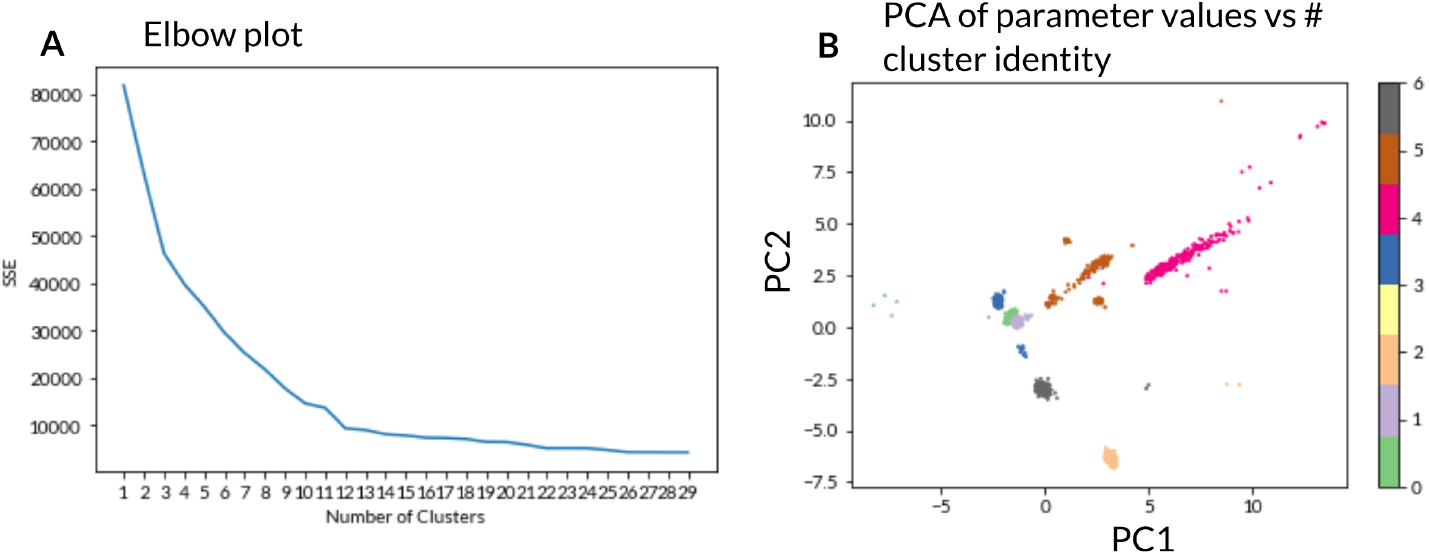
**A**. Elbow plot of SSE vs cluster size. **B**. PCA of parameter values, colour coded by cluster.

Supplementary Movie 1: Visualisation of the AGETs for Wnt and FGF on the cell tracks. AGETs were calculated using the median of the five nearest neighbours.

Supplementary Movie 2: Visualisation of the AGETs for tbxta, tbx16 and tbx6 on the cell tracks. Top panel. AGETs are visualised on the cell tracks in the context of the developing PSM for tbxta (red, left), tbx16 (yellow, middle) and tbx6 (blue, right). Dorsal, up; posterior, left. Bottom panel. Posterior to anterior quantification of AGET expression for the Tbox genes: tbxta (red, left), tbx16 (yellow, middle) and tbx6 (blue, right). X-axis denotes % posterior-anterior position along the developing PSM. Y-axis represents gene expression in arbitrary units. Each dot represents the approximated gene expression for one Tbox gene in a single cell. Curves represent the average expression for each gene along the normalised posterior to anterior axis of the PSM. AGETs were calculated using the median of the five nearest neighbours.

Supplementary Movie 3: Comparison of simulated and approximated Tbox gene expression on the cell tracks. Top panel. Gene expression dynamics simulated using the MAP network are visualised on the cell tracks in the context of the developing PSM for tbxta (red, left), tbx16 (yellow, middle) and tbx6 (blue, right). Dorsal, up; posterior, left. Bottom panel. Posterior to anterior quantification of gene expression for the Tbox genes simulated using the MAP network and compared to the AGETs: tbxta (red, left), tbx16 (yellow, middle) and tbx6 (blue, right). X-axis denotes % posterior-anterior position along the developing PSM. Y-axis represents gene expression in arbitrary units. Each dot represents the simulated gene expression for one Tbox gene in a single cell. Dotted curves represent the average simulated expression for each gene along the normalised posterior to anterior axis of the PSM while solid curves represent the average expression for each gene calculated using the AGETs as in Supplementary Movie 2.

